# A genome assembly of the Atlantic chub mackerel (Scomber colias): a valuable teleost fishing resource

**DOI:** 10.1101/2021.11.19.468211

**Authors:** André M. Machado, André Gomes-dos-Santos, Miguel Fonseca, Rute R. da Fonseca, Ana Veríssimo, Mónica Felício, Ricardo Capela, Nélson Alves, Miguel Santos, Filipe Salvador-Caramelo, Marcos Domingues, Raquel Ruivo, Elsa Froufe, L. Filipe C. Castro

**Affiliations:** CIIMAR – Interdisciplinary Centre of Marine and Environmental Research, U. Porto – University of Porto, Porto, Portugal; Department of Biology, Faculty of Sciences, U. Porto - University of Porto, Portugal; Center for Global Mountain Biodiversity, GLOBE Institute, University of Copenhagen, Copenhagen, Denmark; Center for Macroecology, Evolution, and Climate, GLOBE Institute, University of Copenhagen, Denmark; CIBIO - Centro de Investigação em Biodiversidade e Recursos Genéticos, InBIO - Laboratório Associado, Campus de Vairão, Universidade do Porto, 4485-661 Vairão, Portugal; BIOPOLIS - Program in Genomics, Biodiversity and Land Planning, CIBIO, Campus de Vairão, 4485-661 Vairão, Portugal; Portuguese Institute for the Sea and Atmosphere, I.P. (IPMA), Portugal

**Keywords:** *Scomber colias*, Atlantic chub mackerel, Scombriformes, genome, fisheries management, population dynamics, endocrinology, teleosts

## Abstract

The Atlantic chub mackerel, *Scomber colias* Gmelin, 1789, is a medium-size pelagic fish with substantial importance in the fisheries of the Atlantic Ocean and the Mediterranean Sea. Over the past decade, this species has gained special relevance being one of the main targets of pelagic fisheries in the NE Atlantic. Here, we sequenced and annotated the first high-quality draft genome assembly of *S. colias*, produced with Pacbio HiFi long reads and Illumina Paired-End short reads. The estimated genome size is 814 Mb distributed into 2,028 scaffolds and 2,093 contigs with an N50 length of 4,19 and 3,34 Mb, respectively. We annotated 27,675 protein-coding genes and the BUSCO analyses indicated high completeness, with 97.3 % of the single-copy orthologs in the Actinopterygii library profile. The present genome assembly represents a valuable resource to address the biology and management of this relevant fishery. Finally, this is the fourth high-quality genome assembly within the Order Scombriformes and the first in the genus *Scomber*.

## Data description

### Background and context

The family Scombridae is divided into two subfamilies (Gasterochismatinae and Scombrinae), with 5 genera and around 51 described species, comprising mackerels, bonitos, and tunas [1]. The representative genus of the Scombridae, i.e., *Scomber* includes 4 species: *S. scombrus*, *S. japonicus*, *S. australasicus*, and *S. colias*. The Atlantic chub mackerel, *Scomber colias* Gmelin, 1789, (NCBI Taxonomy ID: 338315) is a small coastal-pelagic fish widely distributed in the Atlantic Ocean from the Bay of Biscay to South Africa (including the Canary, Madeira, Azores, and Saint Helena Islands) and in the Mediterranean Sea [2]. *Scomber colias* is usually found in depths up to 300 m and occupies a key position in the trophic web. This species acts as a link between primary producers and top predators since it feeds mainly on zooplankton and some small pelagic fish and is an essential element of the diet of larger pelagic fish (e.g., tuna, swordfish, and sharks) and marine mammals (e.g., dolphins and seals) [3]. Besides its ecological importance, *S. colias* also supports important commercial fisheries for several countries throughout its distribution range, being an important component in the diet of several local populations [1, 4]. This is probably related to its nutritional value, as this mackerel is a privileged source of important and beneficial fatty acids for human nutrition, particularly Docosahexaenoic acid (DHA), an omega-3 fatty acid [5, 6]. Additionally, *S. colias* is used as bait for the tuna longline and handline fisheries and caught in purse seine and pelagic trawl fisheries which target sardine and/or anchovy [7].

The availability of *S. colias* makes it a sustainable marine resource [6] and a viable alternative to the European’s sardine (*Sardina pilchardus*), which is under fishing restrictions because of population decline. Curiously, fluctuations in the abundance and a northwards shift in the distribution of *S. colias*, with a likely inverse relationship with sardine abundance has been recently demonstrated [8]. Due to its ecological and economic importance, *S. colias* has been the focus of several recent studies focusing on different aspects of its fisheries and biology [3, 8, 9]. Yet, genomic resources for the species are still limited. Only a (liver) transcriptome [10], mitogenome [11], and single-nucleotide polymorphism (SNP) data, obtained through RAD-seq [12], have been described for the species. With the vast majority of the world’s fish stocks already collapsed and with climate change as additional pressure, knowledge of fish genomes is becoming an invaluable tool to address conservation efforts [13, 14]. Here, we report the first high-quality draft genome of *S. colias*, assembled with Illumina and Pacific Biosciences (PacBio) Single-Molecule High-Fidelity (HiFi) reads. This resource provides a critical platform to uncover the species’ adaptive physiological potential in a changing environment. Specifically, it will help understand the current observed populational northward shift, postulated to be part of a more general expansion of species from warmer areas [8]. Moreover, being one of the genomes with higher quality within family Scombridae and the first within the *Scomber* genus, the obtained information will help to improve the conservation, management, and sustainable exploitation of this valuable fish resource as well as of its highly valued congeners.

## Methods

### Sampling and DNA extraction

Two specimens of *S. colias* were collected at two sampling points and time frames. The first specimen was collected in 2017, during the “*Programa Nacional de Amostragem Biológica*” managed by the Instituto Português do Mar e da Atmosfera” (IPMA) in North Atlantic waters (41.501944 N 8.851667 W). From this individual, two tissues were collected and stored in 100% ethanol (muscle) and RNA later (liver). The liver tissue was used to produce and describe the first liver transcriptome of *S. colias* [10]. Muscle tissue was used in the present study, for genomic DNA (gDNA) extraction using the DNeasy Blood and Tissue Kit (Qiagen, Hilden, Germany), following the manufacturer’s instructions. The gDNA was then used for the Illumina Paired-End (PE) sequencing (described below). The second specimen was caught in 2020 near Mira, Portugal (40.5588270 N 9.4529720 W). Immediately upon harvesting, the muscle was snap-frozen in liquid nitrogen. The frozen tissue was shipped to Brigham Young University DNA Sequencing Center (BYU), where gDNA with high molecular weight was extracted from 1.1 g of muscle using the QIAGEN Genomic-tip 20/G Kit. The quality and concentration of gDNA were assessed with Qubit Fluorometric system (ThermoFisher), and the fragment size was determined with a Fragment analyzer (Agilent) before loading on the Sequel II.

### DNA sequencing libraries construction and sequencing

For the first DNA sample, Illumina PE library preparation and sequencing were carried out by Macrogen, Inc (Seoul, Korea), using Illumina HiSeq X Ten platform with 250 bp PE configuration. For the second specimen, PacBio HiFi library preparation and sequencing were performed at BYU, following the manufacturer’s recommendations (Pacific Biosciences) (https://www.pacb.com/wp-content/uploads/Procedure-Checklist-Preparing-HiFi-SMRTbell-Libraries-using-SMRTbell-Express-Template-Prep-Kit-2.0.pdf). The size-selected fraction had 15.3 kb mean read length and was selected on Sage-Elf system (Sage Sciences). The sequencing was conducted on 2 single-molecule, real-time (SMRT) cells using Sequel II system (v.9.0), with a run time of 30h and 2.9 hours pre-extension. The circular consensus analysis was performed in SMRT® Link v9.0 under the default settings (check details at the Suppl. Table 1).

### Raw data quality-control, clean-up, and Genome size estimation

Both short and long read datasets were assessed by FastQC (v.0.11.8) (http://www.bioinformatics.babraham.ac.uk/projects/fastqc/). The Trimmomatic (v.0.38) [15] software was used to filter and remove the low-quality reads as well as the adaptors of the Illumina dataset (LEADING:5 TRAILING:5 SLIDINGWINDOW:4:20 MINLEN:50). Next, trimmed datasets were used to check the overall characteristics of the *S. colias* genome (i.e., genome size, heterozygosity, or unique content), through the GenomeScope2.0 [16]. Briefly, the Jellyfish (v.2.2.10) [17] software was used to build the k-mer frequency distributions, and the final k-mer counts (k-mer 21, 25, 31) were submitted to the GenomeScope2.0 online platform. On the other hand, the HiFi reads were filtered in two ways (Fig. 2). First, the mitochondrial reads were removed by blast searches (blast-n) using a pre-built database of mitochondrial sequences (Database build protocol; 1 – Selection of all complete mitogenomes present in NT-NCBI (nucleotide database of National Center for Biotechnology Information (NCBI)); 2 – Select by taxon (Actinopterygii; Taxonomy ID: 7898); 3 – Sequence length filter 15000-50000 bp; 4 – Build a database with makeblastdb application of ncbi-blast+ (v.2.9.0)). Second, to filter out possible sources of contamination (artefactual or biological), the HiFi reads were checked by blast (blast-n) against NT-NCBI. In the end, only HiFi reads having match hits with more than 90% of identity and query coverage of 50% in Actinopterygii taxon (NCBI Taxonomy ID: 7898) or without match hits at all were considered for further analyses (Fig. 2).

**Figure 1:**
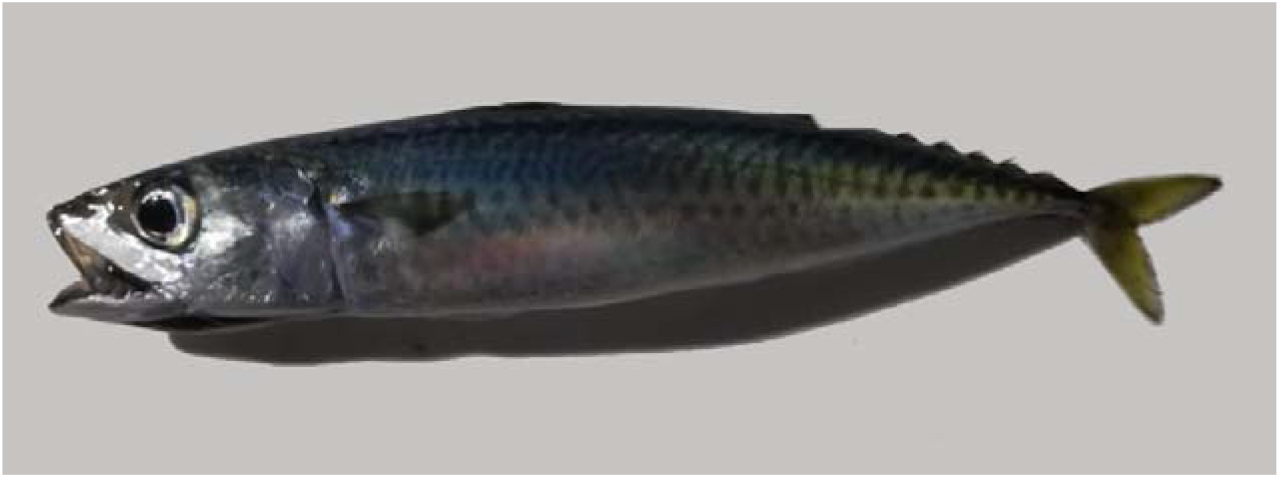
Photograph of Atlantic chub mackerel, *Scomber colias* (Specimen caught in 2020 and used to do the Pacbio HiFi genome assembly).

**Figure 2:**
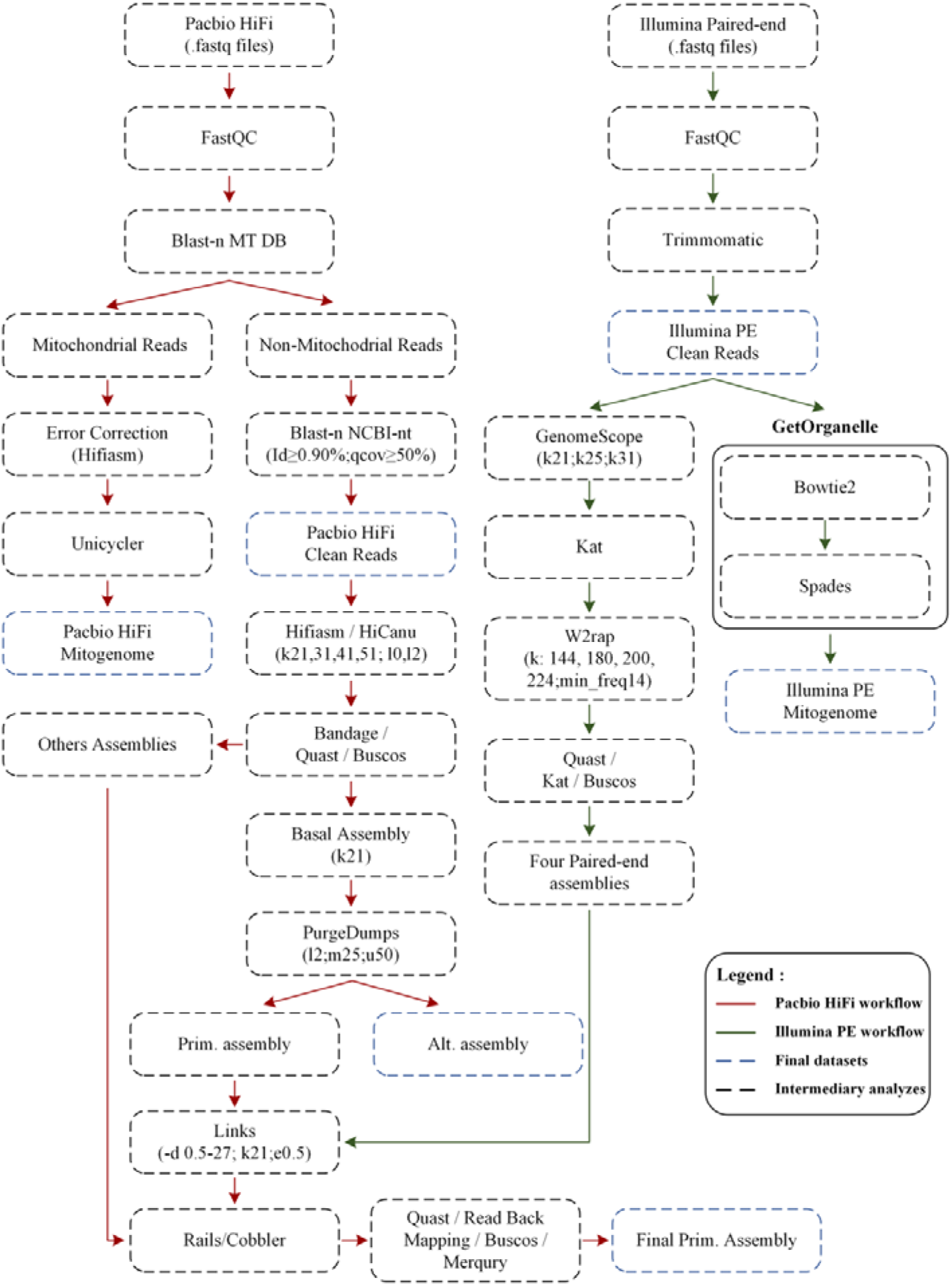
Bioinformatics workflow used to perform the genome assembly of *Scomber Colias* species.

### Mitochondrial genome assembly

Given that two specimens were used for the distinct sequencing approaches, i.e., PacBio HiFi and Illumina PE, the whole mitochondrial genome (mtDNA) was assembled and characterized for both specimens. For specimen one, trimmed Illumina PE reads were used to assemble mtDNA in GetOrganelle (v.1.7.1) [18] (Parameters: - F animal_mt -w 121 -R 10 -k 85,95,105,115,125) (Fig. 2). For specimen two, a new pipeline was designed to produce the mtDNA assembly from the PacBio HiFi long reads (Fig. 2). The PacBio HiFi mtDNA reads, previously filtered (see above), were corrected using Hifiasm (v.0.13-r308) [19] (Parameters: --write-ec). Since Hifiasm is not optimized to assemble circular molecules (expected for the mtDNA), the corrected PacBio HiFi mtDNA reads were assembled using Unicycler (v.0.4.8.) (Parameters: Defaults)[20], a software designed to assemble bacterial genomes and therefore optimized for circular assemblies. Annotation and visual representation of both mtDNA assemblies were produced using MitoZ (v.2.3) [21] (Parameters: -- genetic_code 2; --clade Chordata; --topology circular), using the PE reads for coverage plotting. Furthermore, annotations were manually validated by comparison with other mitochondrial genomes of the genus *Scomber* available at NCBI.

### Nuclear genome assembly and assessment

For whole-genome assembly, a combined approach using short and long-read assemblies was applied (Fig. 2). While the long-read assemblies were mainly used to produce the primary assembly, the short-read assemblies were used to scaffold and improve the contiguity of the basal assembly. In summary, the short-read assemblies were performed with the W2RAP pipeline (v.0.1)[22], following the authors’ protocol (https://github.com/bioinfologics/w2rap). First the Kmer analyses toolkit (KAT) (v.2.4.1) [23] software (hist module) was applied to determine the ideal k-mer cut-off, and then the W2RAP (Parameters: -t 30; -m 500; --min_freq 14; -d 32; --dump_all 1; - k: 144, 180, 200, 224), was used to produce four assemblies (Fig. 2). To generate the long-read assembly, multiple software and parameters were initially tested. The PacBio HiFi reads were assembled in Hifiasm (v.0.13-r308) [19] with different settings (Parameters : k=21, 25, 31, 41, 45, 51, ; l=0, 2) and HiCanu (v.2.1.1) [24] (Parameters: Defaults). While Hifiasm generated 2 pseudo-haplotypes per assembly, the HiCanu generated one merged assembly. To choose the “best” assembly we applied a series of analyses, including Bandage (v.0.8.1) [25] and manual inspection; Benchmarking Universal Single-Copy Orthologs (BUSCO) (v.5.2.2) [26] with Eukaryota and Actinopterygii databases to assess the gene completeness of the assemblies, and Quality Assessment Tool for Genome Assemblies (QUAST) (v.5.0.2) [27] to check the general metrics of the assemblies (Fig. 2). Due to the discrepancies in the length of the Hifiasm primary and alternative pseudo-haplotypes, we opted to concatenate both in a single assembly. At this point, the assembly with the highest complete BUSCO scores, highest contiguity (N50), and longest contig was selected for further analyses. The pseudo-haplotypes were separated by purge_dups (v.1.2.5) [28]. After the first round of purging and inspection by k-mer plot, produced by the KAT tool, the cut-offs were manually adjusted. To assess the influence of purge_dups in the genome, BUSCO (rate of deduplicates) and QUAST (N50 and genomic length per pseudo-haplotype) were used. Next, to improve the contiguity/quality of the assembly, the short read assemblies were used to structurally scaffold the assembly, without the introduction of any new bases in the assembly similarly to [29, 30] (Fig. 2). The four short read assemblies were inputted to the Long Interval Nucleotide K-mer Scaffolder (LINKS) (v.1.8.7) [31] (being used as long reads), and using several distance values, i.e., -d 0.5,1.5,3,9,27 kb, the primary assembly was re-scaffolded interactively for 5 rounds (Additional Parameters: -k 21 -e 0.5). Furthermore, the scaffolded genome and the long-read assemblies, initially produced by Hifiasm and HiCanu and discarded based on contiguity and completeness, were inputted to Cobbler (v.0.6.1) [32] and RAILS (v.1.5.1) [32] pipeline (Parameters: Defaults). This allowed gap-filling of ambiguity regions (produced by short-reads scaffolding) and further re-scaffolding using long read information. To evaluate the final assembly, several metrics and software were used. In addition to BUSCO and QUAST metrics, read back mapping of PE read with Burrows-Wheeler Aligner (BWA) (v.0.7.17-r1198) [33], long reads with Minimap2 (v.2.17) [34] and RNA-Seq with Hisat2 (v.2.2.0) [35, 36] were also applied. To check the consensus quality (QV) and k-mer completeness we used Merqury [37] (v.1.1) (Fig. 2).

### Repeat masking, gene prediction, and annotation

The repetitive elements of the genome were predicted and masked by RepeatMasker (v.4.0.7) [38] using homologous comparisons and *ab initio* predictions. First, the *de novo* library of repetitive elements was created with the RepeatModeler (v.2.0.1) [39]. Next, *ab initio* library, as well as the Dfam_consensus-20170127 [40] and RepBase-20181026 [41], were used in RepeatMaker to softmask the *S. colias* genome assembly.

The genome annotation was performed with the BRAKER2 pipeline (v.2.1.6) [42–44]. Initially, the liver RNAseq reads (Accession number: SRR6367407 [10]) were downloaded, mapped against the *S. colias* genome assembly using Hisat2 (v.2.2.0) [35, 36] (Parameters: Defaults) and converted to bam and sorted files using the SAMtools (v.1.9) [45]. Additionally, we collected 89 proteomes from NCBI RefSeq [46] and Ensembl [47] databases (Suppl. Table 2). Of these, 82 species belong to the Actinopterygii class (32 taxonomic orders), 81 with genome assembly at chromosome level and one at scaffold level. At the date of this genome annotation, only one Scombriforme genome, *Thunnus orientalis*, was annotated (at scaffold level). The remaining seven proteomes were selected from well-established or phylogenetically early-branching species, *Callorhinchus milii*, *Amblyraja radiata*, *Scyliorhinus canicula*, *Lepisosteus oculatus*, *Petromyzon marinus*, *Mus musculus*, *Homo sapiens*. Next, the RNAseq alignment, as well as all the above-mentioned proteomes, were inputted in the BRAKER2 pipeline (Parameters: –etpmode; –softmasking; –UTR=off; –crf; –cores□=30). The final file of predictions (braker.gtf) was further filtered by evidence, keeping only gene predictions with RNAseq or protein evidence (using BRAKER2 auxiliary scripts; selectSupportedSubsets.py), converted to .gff3 format (using the Augustus auxiliary scripts; gtf2gff.pl), and post-processed with Another Gtf/Gff Analysis Toolkit (AGAT) (v.0.6.0). The post-processing stage involved the correction of overlapping gene prediction coordinates and the removal of small or incomplete protein-coding genes (<100 aa; without start/stop codon or both). Furthermore, the proteins were extracted with AGAT tool and functional annotated using InterProScan v.5.44.80 [48] and blast-p searches against RefSeq [46] (Download at 15/05/2021) and SwissProt [49] (Download at 15/05/2021) databases. The homology searches were performed with DIAMOND (v.2.0.11.149) [50] (Parameters: -k 1, -b 10, -e 1e-5, --ultra-sensitive ,--outfmt 6). Finally, the genome as well the annotation datasets were integrated into a website using JBrowse2 [51], a dynamic web platform for genome visualization and analysis that allows easy and interactive exploration of the data provided. The FASTA file containing the genome was indexed with SAMtools Faidx (v.1.9) [45] and added to the JBrowse component, along with the annotation file sorted with “GenomeTools” (v.1.6.1) [52] and indexed with SAMtools Tabix (v.1.9) [53]. In addition to the JBrowse component, ncbi-blast+ (v.2.12.0) [54] was integrated into the webpage allowing the blasted results from the genome, mRNA, protein-coding sequences (CDS), and proteins directly from the website (portugalfishomics.ccimar.up.pt/scombercolias).

### Phylogenomics

To generate a phylogenomic analysis, the proteomes of 15 selected Actinopterygii species, including the Scombriformes species *Thunnus maccoyii* and *T. orientalis*, were downloaded from public databases (Suppl. Table 2). Single-copy orthologs between these 15 species and *S. colias* were retrieved from the protein datasets by constructing protein family clusters using OrthoFinder (v.2.4.0) [55] (Parameters: - M). This resulted in a total of 392 single copy orthologous sequences that were individually aligned using MUSCLE (v.3.8.31) [56] (Parameters: Defaults). Each alignment was trimmed using TrimAl (v.1.2) [57] with a gap threshold of 0.5 (Parameters: -gt 0.5) and afterward concatenated using FASconCAT-G (https://github.com/PatrickKueck/FASconCAT-G). Phylogenetic inferences were conducted in IQ-Tree (v.1.6.12) [58] (Parameters: -bb 10000 -nt AUTO -st AA). The best fitted molecular evolutionary model used in the phylogenetic analyses was JTT+F+R4, which was selected by ModelFinder [59] implemented within IQ-Tree.

### Assessing the Nuclear Receptor and the “*chemical defensome*” repertoire in Scomber colias

To collect the repertoire of the nuclear receptors (NRs) in *S. colias* tblast-n (default parameters) searches were performed in the primary genome assembly. The protein sequences of the DNA binding domains and ligand-binding domains of the *Homo sapiens* NRs were collected from RefSeq [46] database and used as query (NP_000466.2, NP_068804.1, NP_003241.2, XP_005257609.1, NP_001349802.1, NP_068370.1, NP_599022.1, NP_009052.4, NP_001351014.1, XP_005260464.1, NP_002948.1, NP_001257330.1, NP_003288.2, XP_016862607.1, NP_001273031.1, NP_005645.1, NP_001278159.1, NP_004442.3, NP_000167.1, XP_005268879.1, NP_004950.2, NP_201591.2). Next, the regions aligning with the *H. sapiens* sequences were collected, translated to protein using the Bio.Seq module of biopython (v.1.75) [60], and blasted (blast-p) against a local database containing the NRs proteins of *Danio rerio* (*D. rerio* NRs Database protocol; 1 – NRs sequences and classifications were retrieved from [61]; 2 – The NRs database was build using the makeblastdb application of ncbi-blast+ (v.2.12.0)). For each NRs sequence in *S. colias,* the best blast hit in the *D. rerio* database was collected. In some cases, several nuclear receptors of *S. colias* matched the same receptor in *D. rerio*. In these cases, the nucleotide sequences of *S. colias* were re-validated against the NT-NCBI database, and all sequences matching different GeneID’s in the same organism were kept in the final table of NRs. In parallel, and to assess the genome annotation performed by BRAKER2, the genomic coordinates of the regions aligning to *H. sapiens* were searched and identified in the annotation files.

To identify the genes related to the “*chemical defensome*”, target genes were selected based on a previous report profiling the “*chemical defensome*” of teleost species [62]. Next, gene names were used as queries to search the deduced *S. colias* genome annotation, a simple but successful approach for well-annotated genomes such as *D. rerio* [62]. When gene names were not retrieved from *S. colias* genome annotation (i.e. *fthl*, *gstp*, *hsph*, *maff*, *nme8*, *slc21*), further tblast-n searches were performed (default parameters, except -max_hsps 1 to keep the best query-subject pair) in the primary genome assembly, using *D. rerio* sequences as a query.

### Demography with pairwise sequentially Markovian coalescent (PSMC)

To explore the variation in the demographic history of the species, the pairwise sequentially Markovian coalescent (PSMC) strategy was applied [63], following the authors’ instructions (https://github.com/lh3/psmc). Briefly, the PE short reads were aligned to the repeated masked genome assembly using BWA (v.0.7.17-r1198) (Parameters: bwa mem) [33], and the output converted to bam and sorted using SAMtools (v.1.9) [45] (Function: Sort; Parameter: Default). Next, Picard Tools (v.2.19.2) (http://broadinstitute.github.io/picard/) was used to remove duplicate reads (Function: MarkDuplicates; Parameters: Default), and SAMtools used for mapping quality filtering and SNP calling (Function: mpileup; Parameters: -Q 30 -q 30 -C 50). The BCFtools (v.1.9) was applied to extract consensus sequences (Function: call; Parameters: -c) and the subscript vcfutils (from SAMtools) for filtering the output for a minimum depth of 25, a maximum depth of 150, and a min RMS mapQ of 20 (Function: vcf2fq; Parameters: -d 25 -D 150 -Q 20). The resulting fastq file was converted to a PSMC compatible input format using fq2psmcfa with a minimum quality threshold of 20 (Parameters: -q 20). Inferences of population history were performed by running PSMC for 25 iterations (Parameters: -N 15, -r 5, -p 4*4□+□13*2□+□4*4□+□6) following the recent PSCM estimations on Scombriformes [64]. Furthermore, to account for uncertainties in the PSMC estimates, bootstrapping of 100 replicates was performed using the splitfa script provided by the PSMC authors (https://github.com/lh3/psmc). Finally, to scale the demographic estimations, a mutation rate (μ) of 7.3 × 10^−9^ substitutions/site/generation was used, based on the recent estimation for the Scombriformes species *Thunnus albacares* [64], and a generation time for *S. colias* of 2 years [7, 65].

### Data Validation

To produce the *S. colias* genome assembly two sequencing strategies were used, i.e., Illumina PE short reads and PacBio HiFi long reads. The PE dataset was used to assess the genomic proprieties of the *S. colias* species and scaffold the long-read assembly, while the HiFi reads were used to perform the primary genome assembly and the gap closing (Fig. 2).

The Illumina sequencing yielded 149 M of PE reads and the PacBio sequencing generated 1,7 M of HiFi reads (Table 1). Trimmed short reads were used to estimate the genome size (817 Mb), heterozygosity rate (1.31%), and genome repeat content (approximately 26%) using GenomeScope2 (Fig. 3, Suppl. Table 2). In parallel, the HiFi dataset was inspected and mitochondrial reads, as well as possible sources of contamination, were removed (0,31% of the initial dataset) (Table. 1).

**Table 1.** General statistics of read datasets used to perform the *Scomber colias* genome assembly.

| Sample | Sequencing Type | Library type | Platform | Insert size (bp) | Number of Reads (Before clean-up) | Number of Reads (After clean-up) | Application |
| --- | --- | --- | --- | --- | --- | --- | --- |
| Sco_PH | WGS | Long reads | PacBio Sequel II System | 15500 | 1,792,104 | 1,786,541 | Genome Assembly, Gap Closing, Assessment |
| Sco_PE | WGS | Short reads | HiSeq X Ten | 478 | 149,564,893 | 84,738,393 | Scaffold, Assessment |

**Figure 3:**
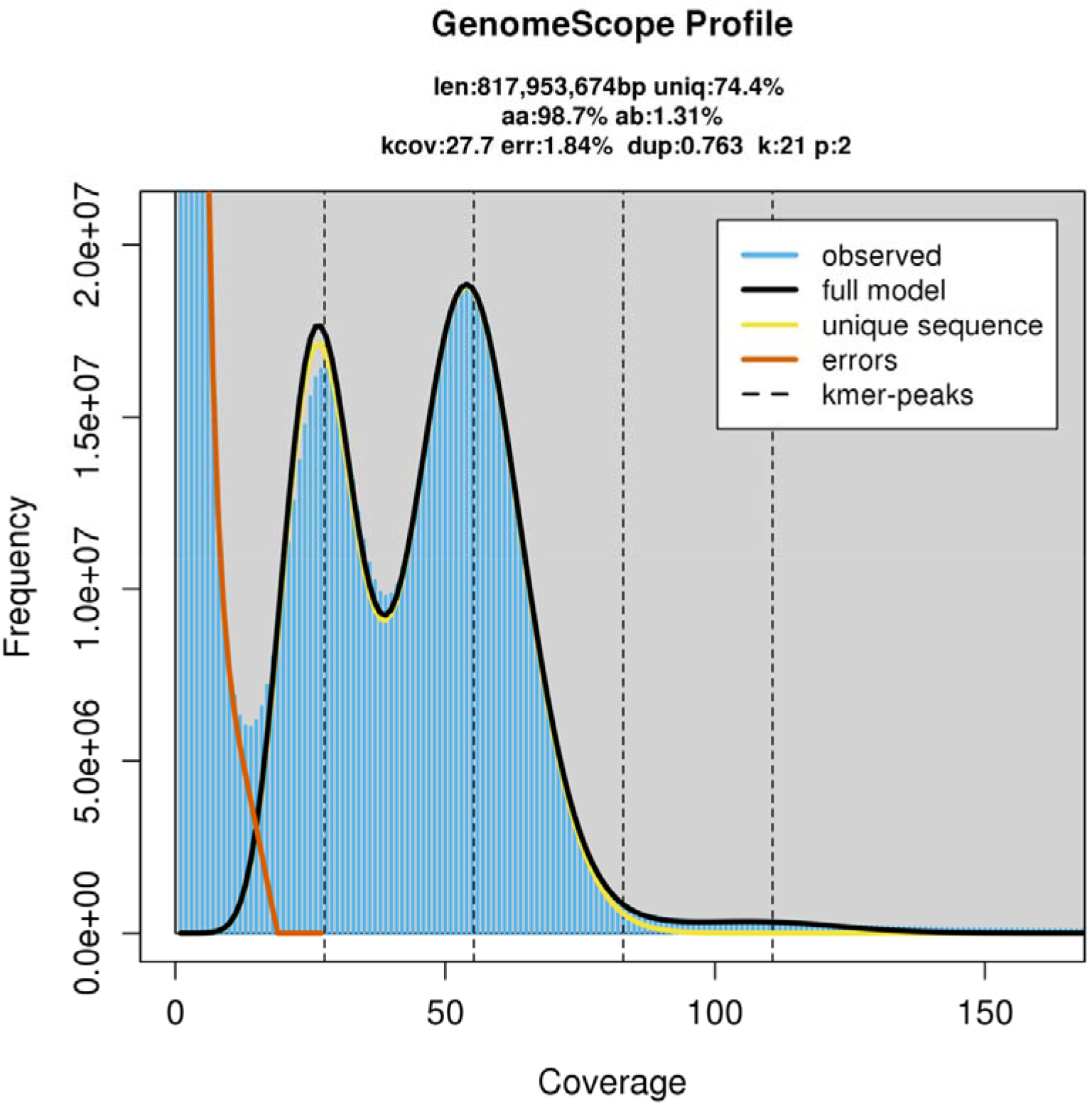
Genomescope2 plot with K-mer spectra content and fitted models of the *Scomber colias* Illumina PE dataset.

For the mtDNA assemblies, a total of 38,868 mtDNA PE reads were filtered by GetOrganelle and a total of 792 mtDNA PacBio HiFi reads were filtered by blast-n search. The two assemblies had the same length of 16,570 bp and differed from each other 0.29% (uncorrected *p*-distances). Furthermore, the PE and PacBio HiFi mtDNA assemblies differed from the *S. colias* mtDNA assembly available on NCBI (Accession number: AB488406.1 [11]), by 0.35% and 0.40% (uncorrected *p*-distances), respectively. The mtDNA gene content and arrangement is expected for most fishes and the standard for vertebrates [66], consisting of 13 protein-coding genes, 22 transfer RNA (trn), and two ribosomal RNA (rrn) (Fig. 4).

**Figure 4:**
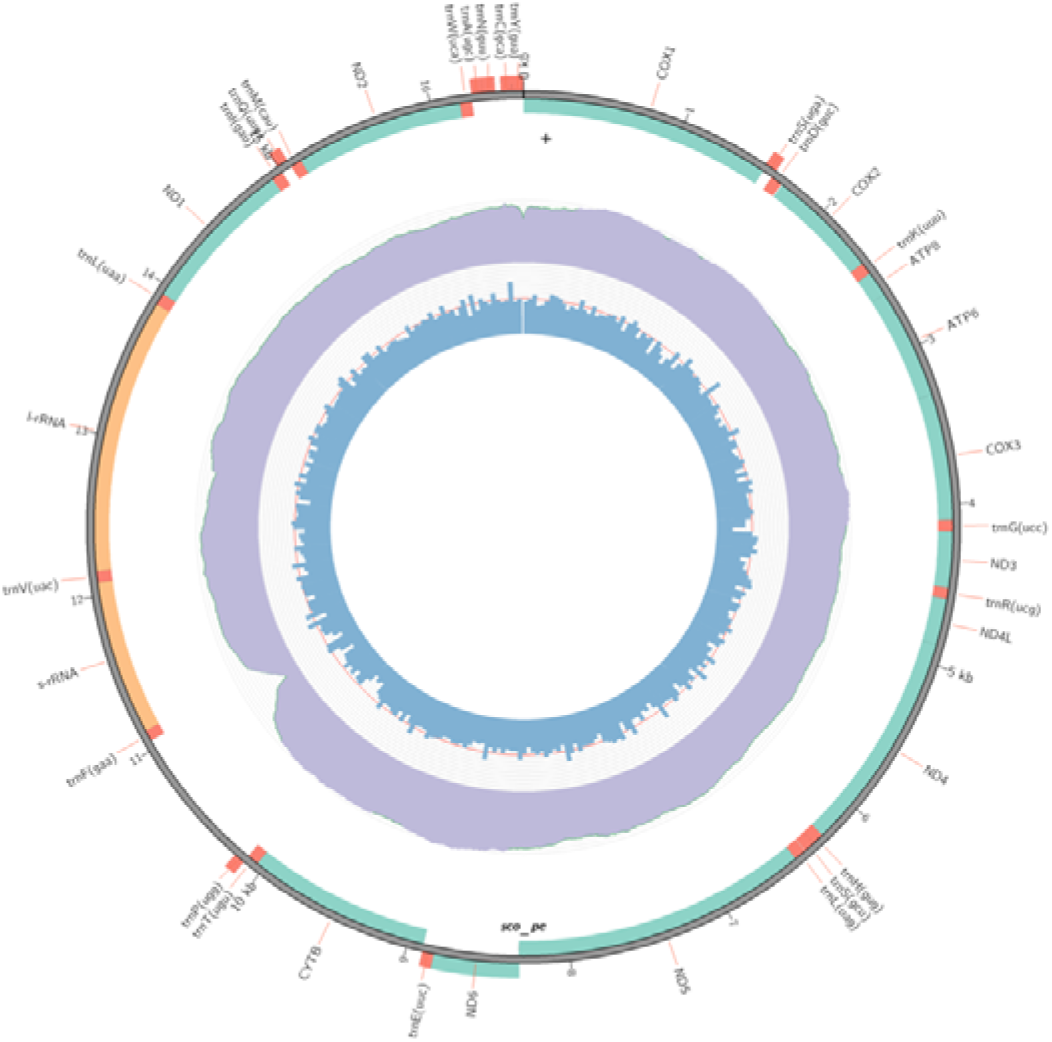
Circular mitochondrial genome assembly of *Scomber colias*, obtained from the Illumina PE dataset (equal to that obtained from the PacBio HiFi long reads assembly). From the center to the outmost features: the GC content distribution; sequencing depth distribution of aligned Paired-End reads; and gene elements (i.e, PCGs, rRNA genes, tRNA genes)

The primary genome assembly was produced using the filtered PacBio HiFi reads and a combination of several software/settings. Following the above-mentioned criteria (Material and Methods – Nuclear genome assembly and assessment) the Sco_k21 assembly was selected (statistics in Suppl. Table 4), both pseudo-haplotypes merged, and subject to purge_dups. Although the purge_dumps generated a primary and an alternative assembly, only the primary was used in the next steps. At the same time, four short-read genome assemblies were performed with the W2RAP software, and the contigs with more than 500bp were used as “long reads” to scaffold the primary assembly (please consult statistics in the Suppl. Table 5). Importantly, during the scaffolding process, only the structural information of short-read assembly was used, without the inclusion of any base. Lastly, the remaining non-basal long read assemblies were used to fill the gaps inserted during the scaffolding stage. The final assembly (primary assembly) of *S. colias* yields a genome size of 814 Mbp, distributed in 2,028 scaffolds and 2,093 contigs with an N50 length of 4,19 and 3,34 Mbp, respectively. On the other hand, the alternative assembly had 807 Mbp and 5,908 contigs with an N50 length of 0.47 Mbp (Table 2). The BUSCO analyses, at the nucleotide level, in Eukaryota and Actinopterygii datasets showed high levels of completeness for both primary (96.9% and 97.3% of single-copy orthologs) and alternative assemblies (93.3% and 96 % single-copy orthologs) (Table 2). Consistently, Merqury determined high QV (Primary – 56.53 %; Alternative – 54.99%) and k-mer completeness (Primary – 86.11%; Alternative – 84.60%) values for both assemblies (Table 2). In the primary assembly, the k-mer analyses (via Merqury) showed a low level of k-mer duplication in the genome (color blue, green purple, and orange in Fig. 5a), indicating a high level of haplotype uniqueness (color red Fig. 5a), and a similar k-mer distribution pattern to GenomeScope2 (performed with Illumina PE reads). Additionally, we found a high rate of mappings of the Illumina, PacBio, and RNA-Seq reads, against the primary assembly of 95 %, 99.8 %, and 90.02 %, respectively. Overall, these results provide evidence of the high quality of the *S. colias* genome assembly (Table 2).

**Table 2.**
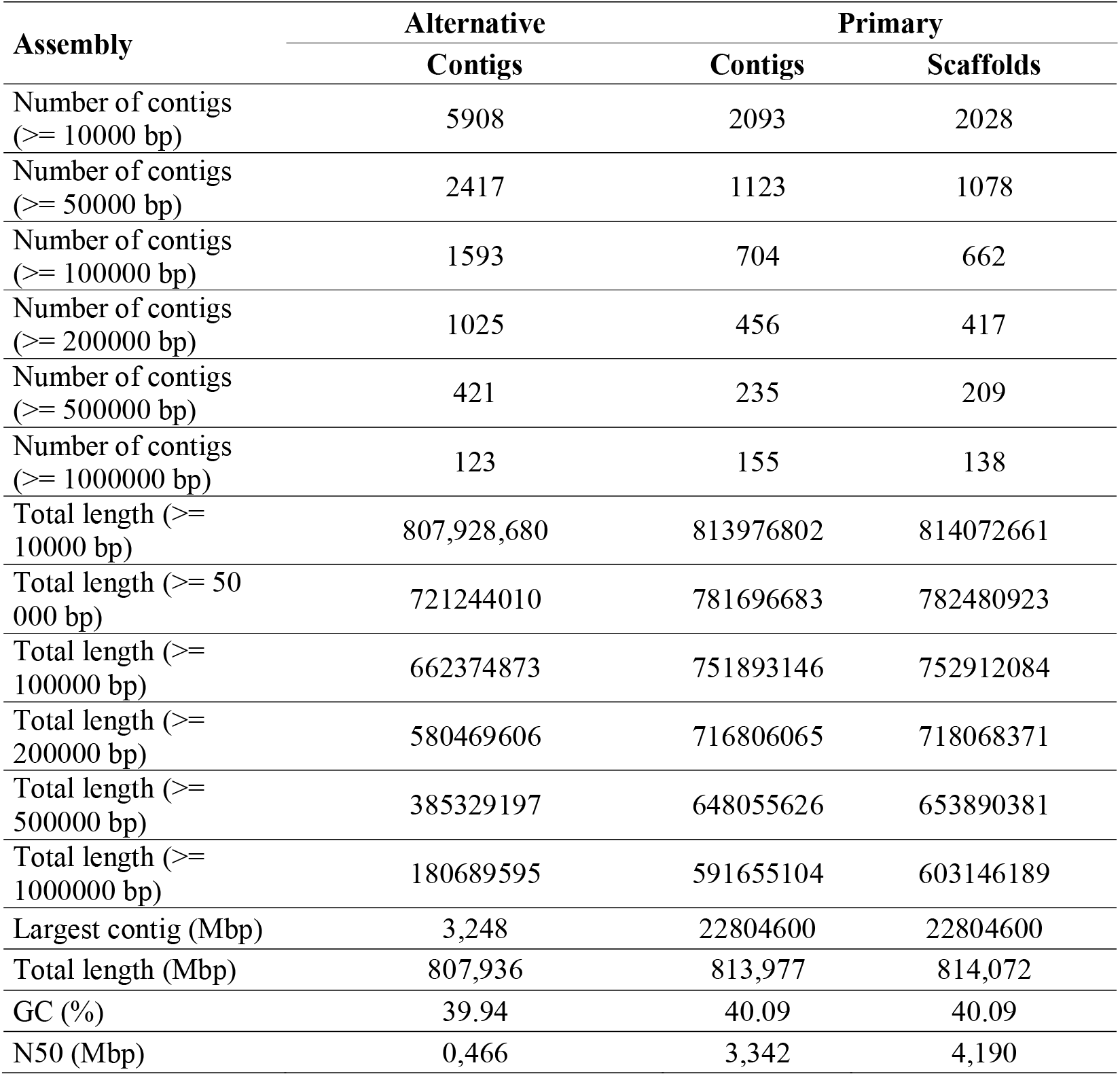
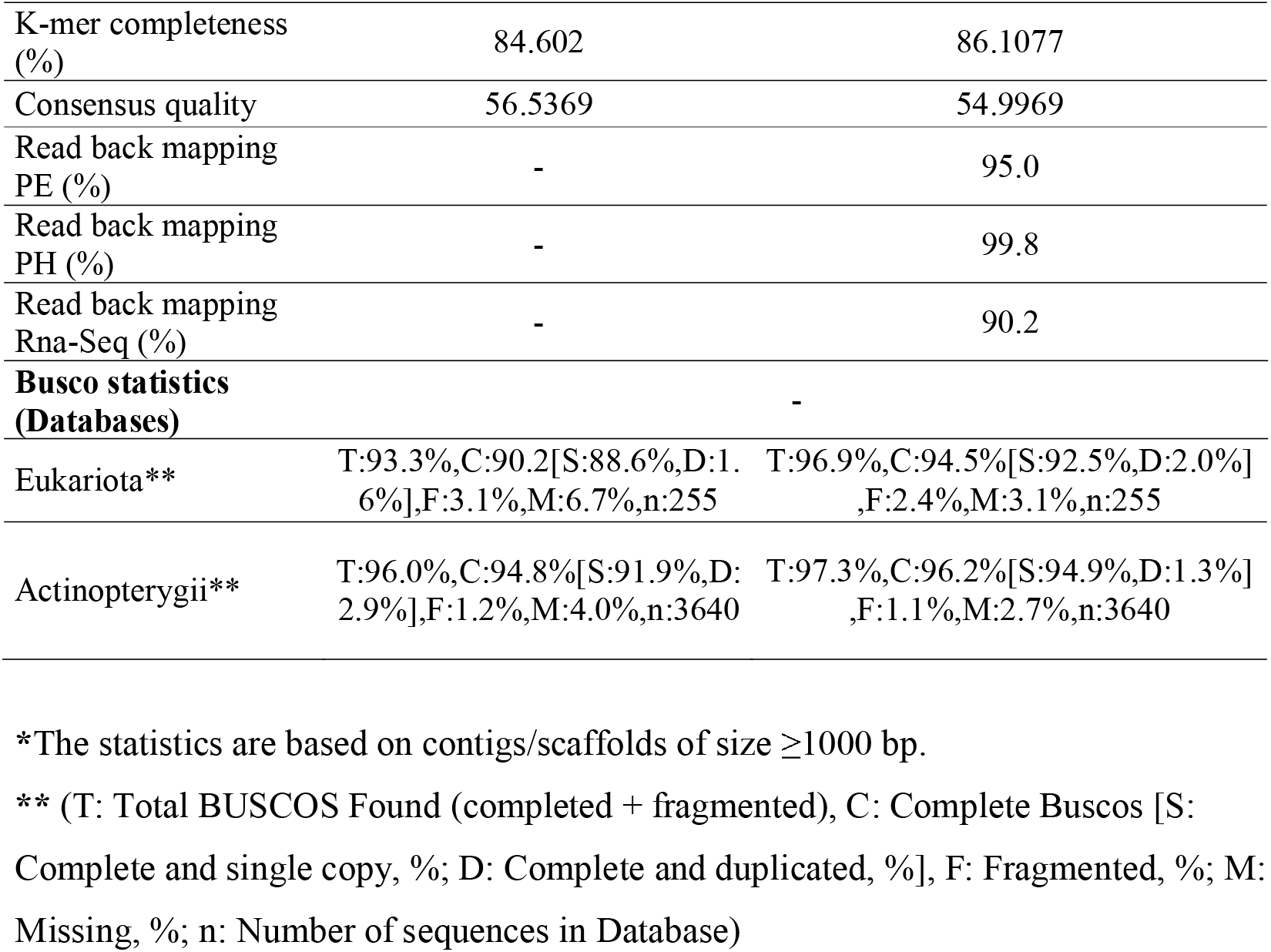
Statistics of *Scomber colias* genome assembly.

| Assembly | Alternative | Primary |  |
| --- | --- | --- | --- |
|  | Contigs | Contigs | Scaffolds |
| Number of contigs (>= 10000 bp) | 5908 | 2093 | 2028 |
| Number of contigs (>= 50000 bp) | 2417 | 1123 | 1078 |
| Number of contigs (>= 100000 bp) | 1593 | 704 | 662 |
| Number of contigs (>= 200000 bp) | 1025 | 456 | 417 |
| Number of contigs (>= 500000 bp) | 421 | 235 | 209 |
| Number of contigs (>= 1000000 bp) | 123 | 155 | 138 |
| Total length (>= 10000 bp) | 807,928,680 | 813976802 | 814072661 |
| Total length (>= 50 000 bp) | 721244010 | 781696683 | 782480923 |
| Total length (>= 100000 bp) | 662374873 | 751893146 | 752912084 |
| Total length (>= 200000 bp) | 580469606 | 716806065 | 718068371 |
| Total length (>= 500000 bp) | 385329197 | 648055626 | 653890381 |
| Total length (>= 1000000 bp) | 180689595 | 591655104 | 603146189 |
| Largest contig (Mbp) | 3,248 | 22804600 | 22804600 |
| Total length (Mbp) | 807,936 | 813,977 | 814,072 |
| GC (%) | 39.94 | 40.09 | 40.09 |
| N50 (Mbp) | 0,466 | 3,342 | 4,190 |

|  |  |  |
| --- | --- | --- |
| K-mer completeness (%) | 84.602 | 86.1077 |
| Consensus quality | 56.5369 | 54.9969 |
| Read back mapping PE (%) | - | 95.0 |
| Read back mapping PH (%) | - | 99.8 |
| Read back mapping Rna-Seq (%) | - | 90.2 |
| **Busco statistics (Databases)** | - |  |
| Eukariota** | T:93.3%,C:90.2[S:88.6%,D:1.6%],F:3.1%,M:6.7%,n:255 | T:96.9%,C:94.5%[S:92.5%,D:2.0%],F:2.4%,M:3.1%,n:255 |
| Actinopterygii** | T:96.0%,C:94.8%[S:91.9%,D:2.9%],F:1.2%,M:4.0%,n:3640 | T:97.3%,C:96.2%[S:94.9%,D:1.3%],F:1.1%,M:2.7%,n:3640 |

\*The statistics are based on contigs/scaffolds of size $\geq 1000$ bp.
\*\* (T: Total BUSCOS Found (completed + fragmented), C: Complete Buscos [S: Complete and single copy, %; D: Complete and duplicated, %], F: Fragmented, %; M: Missing, %; n: Number of sequences in Database)

**Figure 5:**
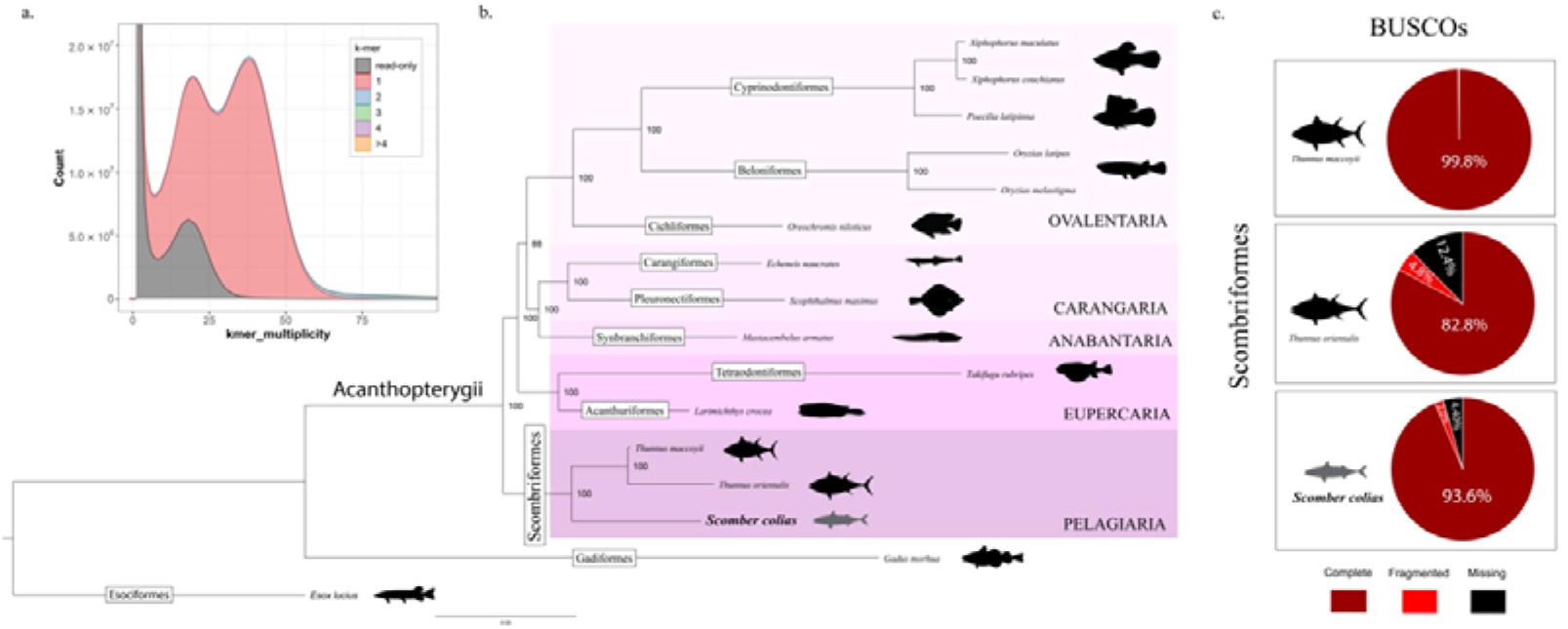
Validation of the genome assembly and annotation process. a. K-mer analyses of the *Scomber colias* genome assembly (Merqury). b. Maximum Likelihood phylogenetic tree based on the concatenated alignments of amino acid sequences of 392 single-copy orthologs retrieved by OrthoFinder. Bootstrap values are shown next to the nodes. c. BUSCOs scores were obtained from searching the proteomes of the three Scombriformes species with genome annotation available, against the actinopterygii_odb10 (n:3640) lineage.

The RepeatMasker software masked 29.62% of the bases in the primary genome assembly. The major part of the masked regions was linked to DNA elements (11.66%), long interspersed nuclear elements (4.11%), long terminal repeats (2.58%), and simple repeats (2.88%). Furthermore, 8.62 % of the genome was masked and annotated as Unclassified, and only a small percentage was classified as short interspersed nuclear elements, Small RNA or Satellites repeats (Table 3). The genome annotation process generated about 27,675 protein-coding genes and 30,999 protein-coding sequences. On average, we found 9,5 exons and 1,656 bp of length per CDS (Table 4). 30,355 of the CDS had at least one blast-p hit in Swissprot and/or RefSeq databases, 27,101 were identified in the InterPro database and 21,664 of these were classified as belonging to a specific homolog superfamily (Table 5).

**Table 3.**
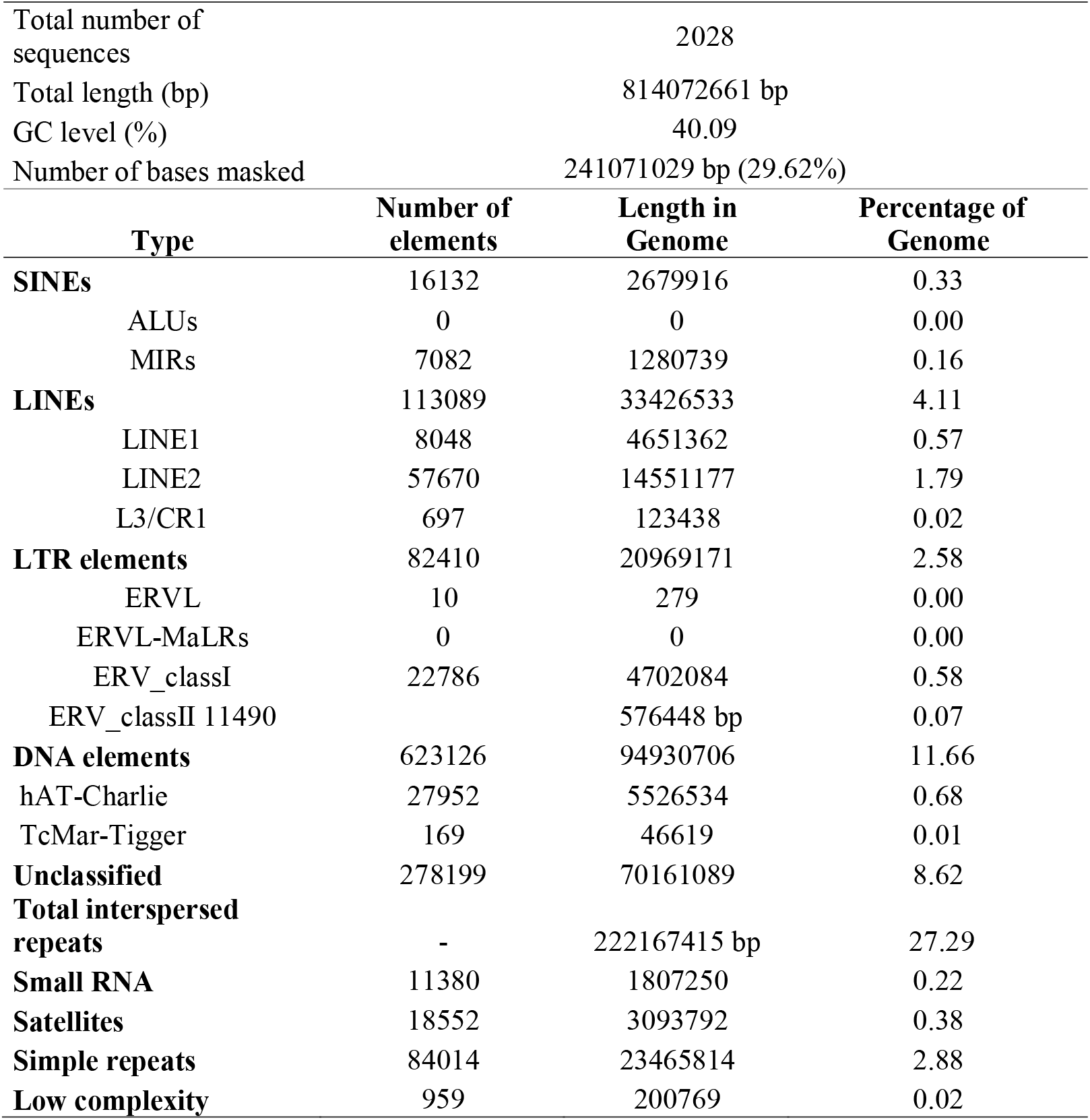
Report of RepeatMasker software. This report contains the statistics of the repetitive elements in *Scomber colias* genome assembly.

**Table 4.**
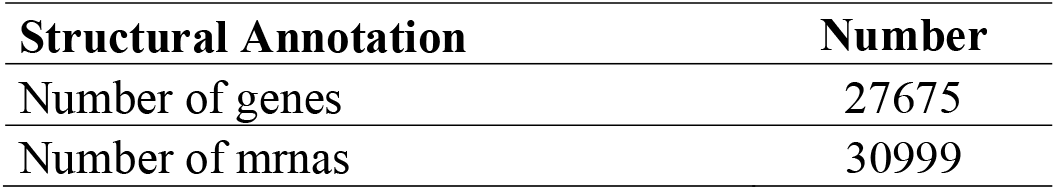
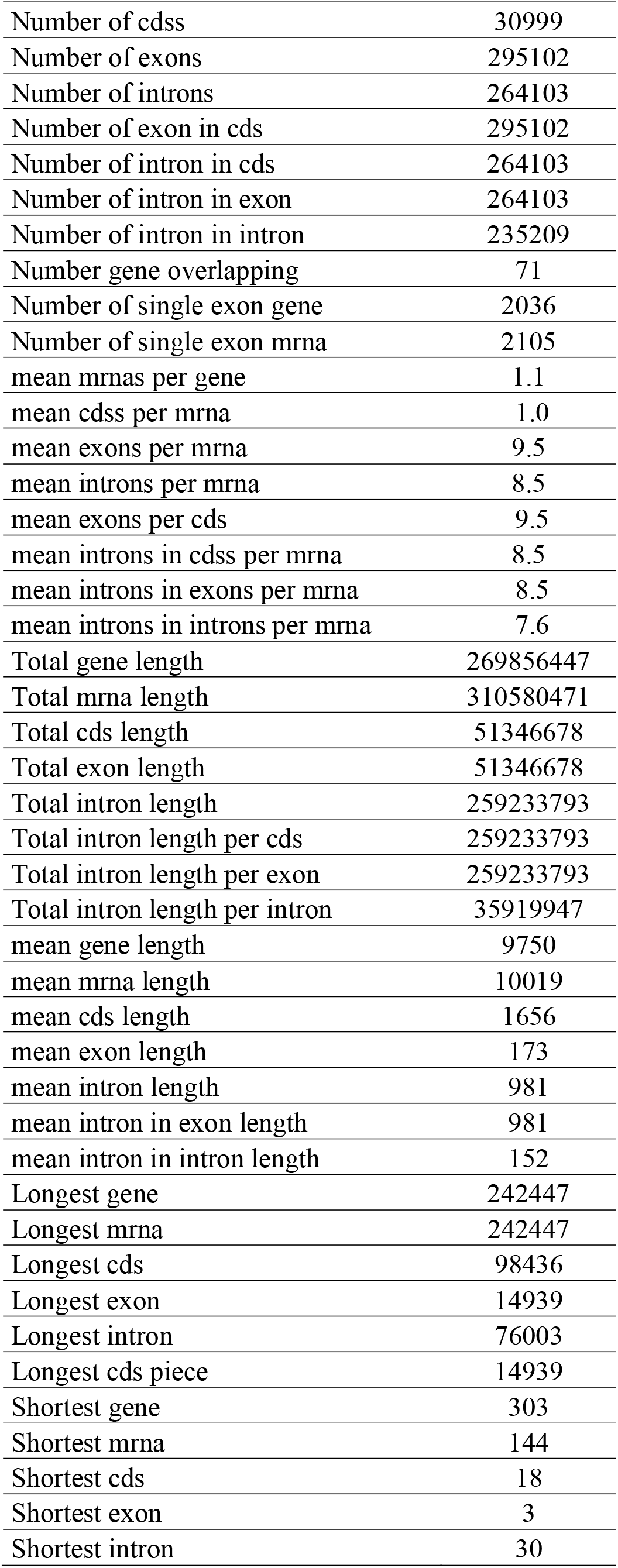
Structural annotation report of *Scomber colias* genome assembly.

| Structural Annotation | Number |
| --- | --- |
| Number of genes | 27675 |
| Number of mrnas | 30999 |
| Number of cdss | 30999 |
| Number of exons | 295102 |
| Number of introns | 264103 |
| Number of exon in cds | 295102 |
| Number of intron in cds | 264103 |
| Number of intron in exon | 264103 |
| Number of intron in intron | 235209 |
| Number gene overlapping | 71 |
| Number of single exon gene | 2036 |
| Number of single exon mrna | 2105 |
| mean mrnas per gene | 1.1 |
| mean cdss per mrna | 1.0 |
| mean exons per mrna | 9.5 |
| mean introns per mrna | 8.5 |
| mean exons per cds | 9.5 |
| mean introns in cdss per mrna | 8.5 |
| mean introns in exons per mrna | 8.5 |
| mean introns in introns per mrna | 7.6 |
| Total gene length | 269856447 |
| Total mrna length | 310580471 |
| Total cds length | 51346678 |
| Total exon length | 51346678 |
| Total intron length | 259233793 |
| Total intron length per cds | 259233793 |
| Total intron length per exon | 259233793 |
| Total intron length per intron | 35919947 |
| mean gene length | 9750 |
| mean mrna length | 10019 |
| mean cds length | 1656 |
| mean exon length | 173 |
| mean intron length | 981 |
| mean intron in exon length | 981 |
| mean intron in intron length | 152 |
| Longest gene | 242447 |
| Longest mrna | 242447 |
| Longest cds | 98436 |
| Longest exon | 14939 |
| Longest intron | 76003 |
| Longest cds piece | 14939 |
| Shortest gene | 303 |
| Shortest mrna | 144 |
| Shortest cds | 18 |
| Shortest exon | 3 |
| Shortest intron | 30 |

**Table 5.** Functional annotation report of *S. colias* genome assembly.

| Functional Annotation | Number |
| --- | --- |
| Swissprot / RefSeq | 30355 |
| InterPro | 27101 |
| CDD | 12832 |
| Coils | 7705 |
| GO | 18643 |
| Gene3D | 22209 |
| Hamap | 463 |
| KEGG | 1402 |
| MetaCyc | 1140 |
| MobiDBlite | 16765 |
| PIRSF | 1755 |
| PRINTS | 7143 |
| Pfam | 25708 |
| ProSitePatterns | 8082 |
| PrositeProfiles | 16229 |
| Reactome | 7376 |
| SFDL | 114 |
| SMART | 14906 |
| SUPERFAMILY | 21664 |
| TIGRFAM | 1427 |

To validate the protein-coding sequences we performed phylogenetic (via OrthoFinder) and BUSCO analyses (using Actinopterygii library profile) (Fig. 4 b, c). Of the 16 Actinopterygii proteins datasets imputed to OrthoFinder, 98.3% were assigned to 29,066 orthogroups, with 12,334 orthogroups present in all species (Suppl. Table 6). Furthermore, a total of 392 single-copy orthologues were retrieved by OrthoFinder and used for the phylogenomic analysis. Alignment, trimming and concatenation of all single-copy orthologues, resulted in a final 120,886 aa long supermatrix alignment that was used for the phylogenomic inference in IQ-Tree. The resulting Maximum Likelihood phylogenetic tree has maximum support for almost all nodes (Fig. 5b). The phylogeny recovered the reciprocal monophyletic Acanthopterygii groups Pelagiaria, Eupercaria, Anabantaria, Carangaria, and Ovalentaria, with Pelagiaria being the basal clade and represented by the three Scombrifomes, including *S. colias* (Fig. 5b). These results are in accordance with the most recent phylogenomic study of ray-finned fishes [67], as well as the Ensembl Compara Species Tree (https://m.ensembl.org/info/about/speciestree.html). The BUSCOs analysis showed the *S. colias* proteome with 93.6% of the groups complete, 2% fragmented, and 4.4% missing (Fig. 5c). In comparison, *T. maccoyii* had 99,8 % BUSCO groups complete while *T. orientalis* had only 82,8 %. These results are expected, since the *T. maccoyii* genome assembly, part of the Vertebrate Genome Project [68], was built at chromosome level with multiple technologies (including 46x PacBio data, 46x 10X Genomics Chromium data, BioNano data, and Arima Hi-C data) and several manual curation steps [69]. On the other hand, both *T. orientalis* [70] and *S. colias* were built at scaffold level using only short and long read information.

We further explored the quality of the annotation by investigating the repertoire of the nuclear receptor (NR) superfamily in the *S. colias* assembly. NRs are critical molecular physiology components, with vital roles in animal physiology and disruption [71]. Moreover, their exact NR gene complement in vertebrate lineages has been shown to vary [61]. We were able to deduce the existence of 76 NRs in the *S. colias* genome (Suppl. Table 7), in line with the repertoire described for other teleost species [72]. Among the retrieved NRs we found those that are key components of the “*chemical defensome*”—an ensemble of related and unrelated genes that protect organisms against chemical stressors, and thus critical under anthropogenic chemical build-up and climate change scenarios —such as the xenobiotic-inducible pregnane X receptor (*pxr*, *nr1i2*) [62, 73]. Subsequent analysis, using gene names, further suggested the presence of gene annotations for the vast majority of the reported members of the teleost “*chemical defensome*” in *S. colias*, similarly to that described for D. rerio [62]. Additional blast searches were performed for a reduced set of genes (*fthl*, *gstp*, *hsph*, *maff*, *nme8,* and *slc21*), uncovering possible homologs for this gene subset, except for a single member of the GST family (*gstp*) (Suppl. Table 8).

We additionally validated our dataset by examining the present population structure of the species, since the genome may also provide clues regarding its past demographic history [63]. One popular method to produce these inferences is the pairwise sequentially Markovian coalescent (PSMC) model, here implemented to the *S. colias* final genome assembly. Since PSMC requires an estimation of the genome-wide mutation rate, and since this has never been produced for *S. colias*, we used the recently estimated genome-wide mutation rate of the yellowfin tuna, *T. albacares* of 7.3 × 10^−9^ mutations/site/generation [64]. The results suggest a past population expansion between 160,000 – 115,000 years ago with maximum effective population size (N_e_) of 36,000, during the end of the Mid-Pleistocene Transaction, corresponding to the Eemian (i.e., the last interglacial period) and the transition between the Marine Isotope Stages (MIS) 6 to 5, (Fig. 6). This population expansion is followed by an apparent decrease in the N_e_ to around 25.000, at the beginning of the Late Pleistocene, corresponding to the entering of the Last Glacial Period. These results, suggesting the influence of the climatic changes from the Pleistocene glaciation cycles on the N_e_, are following other recent studies on Scombriformes, e.g., Pacific Sierra mackerel, *Scomberomorus sierra* [74], and the Indo-Pacific Yellowfin Tuna *T. albacares* [64], as well as in other pelagic marine species, e.g., the killer whale [75].

**Figure 6:**
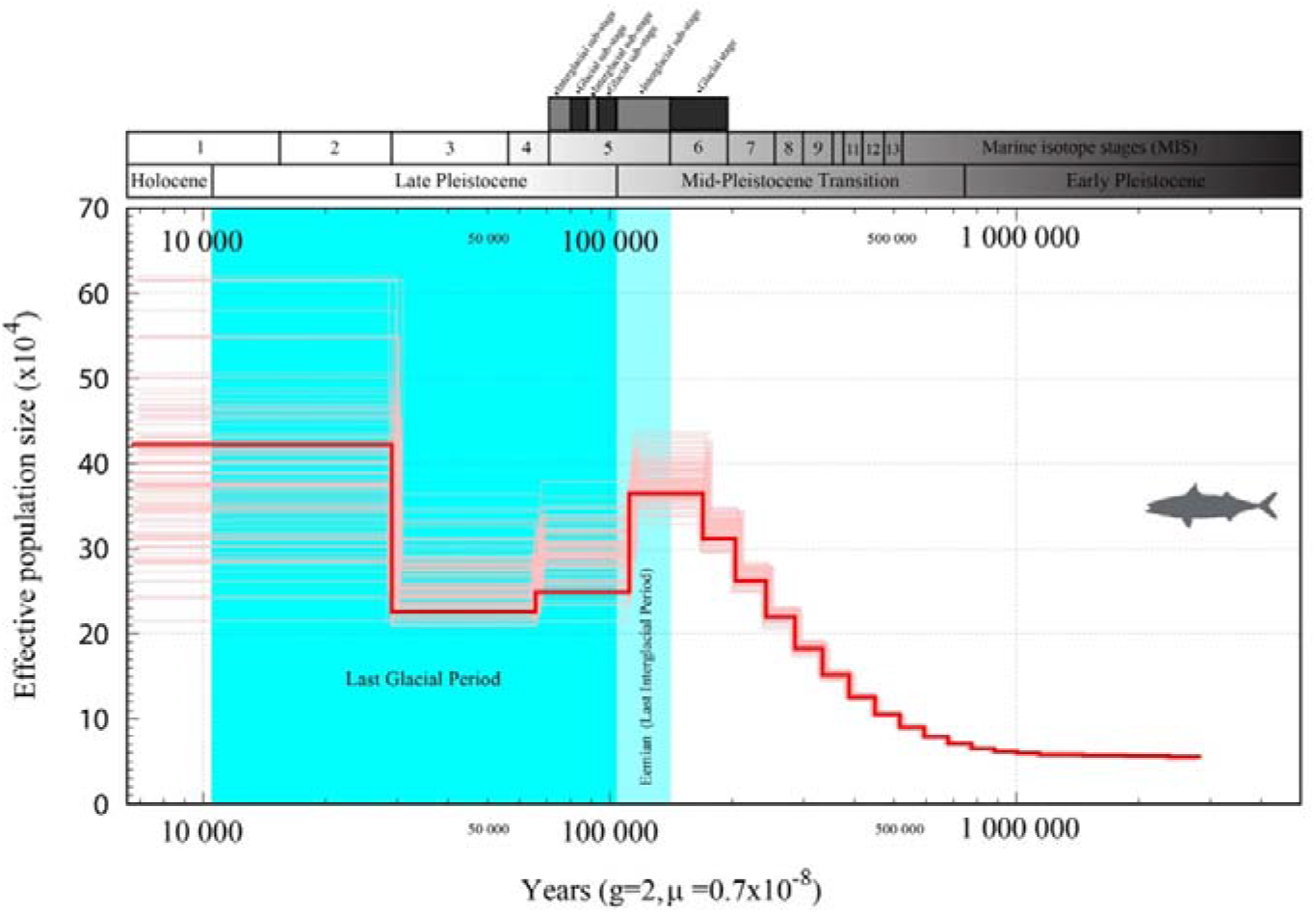
Pairwise sequentially Markovian coalescent (PSMC) estimates from the *Scomber colias* genome assembly. Estimations were obtained using a generation time of 2 years and genome-wide mutation rate of 7.3 × 10^−9^ mutations/site/generation. Effective population size (N_e_) is presented in the left vertical axis and changes estimated since the present over the last 3 myr in the horizontal axis.

### Reuse potential

This study reports the first genome assembly of Atlantic chub mackerel. *Scomber colias* is a valuable marine resource, with a high impact on the fisheries of several countries on the west coast of the Atlantic Ocean and/or the Mediterranean Sea. Ecologically, this species establishes an important link between primary producers and top predators of the marine trophic web. Despite the ecological and economic importance of *S. colias*, few genomic resources are available for this species. Thus, this genome is timely and is expected to contribute to the effective conservation, management, and sustainable exploitation of *S. colias* species in the Anthropocene. Additionally, this genome will be a key tool to decipher biological features of *S. colias*, such as population dynamics, physiology, or endocrinology.

### Data Records

The raw datasets of PacBio HiFi and Illumina sequencing were deposited in the NCBI Sequences Read Archive under the Bioproject: PRJNA769550. Additionally, both primary and alternative pseudo-haplotype assemblies were submitted to NCBI GenBank (Accession number: JAJDFG000000000 and JAJDFH000000000). The mitochondrial genome assemblies and annotations were submitted to GenBank (Accession number: OK501306 and OK501307). The four W2RAP assemblies as well as the genome annotation files were uploaded to figshare online repository (https://doi.org/10.6084/m9.figshare.17025506.v1). The genome and annotation datasets also can be interactively explored at the website - http://portugalfishomics.ciimar.up.pt/app/scombercolias/.

## Declarations

## List of abbreviations

AGAT: Another Gtf/Gff Analysis Toolkit
BUSCO: Benchmarking Universal Single-Copy Ortholog
BWA: Burrows-Wheeler Aligner
CDS: Coding sequences
DHA: Docosahexaenoic acid
Gbp: gigabase pair(s)
gDNA: Genomic DNA
KAT: Kmer analyses toolkit
LINKS: Long Interval Nucleotide K-mer Scaffolder
Mbp: megabase pair(s)
mtDNA: Mitochondrial genome
NCBI: National Center for Biotechnology Information
NRs: Nuclear receptors
NT-NCBI: Nucleotide database of NCBI
PacBio: Pacific Biosciences
PE: Paired-end
PSMC: Pairwise sequentially Markovian coalescent
QUAST: Quality Assessment Tool for Genome Assemblies
QV: consensus quality value
SMRT: Single Molecule, Real-Time
SNP: Single-nucleotide polymorphism

## Ethical approval

This work has been approved by the CIIMAR ethical committee and by CIIMAR Managing Animal Welfare Body (ORBEA) according to the European Union Directive 2010/63/EU.

## Author contributions

LFCC designed and conceived this work; MF, RC, and NA collected the samples; AMM, AGS, EF, LFCC wrote the manuscript; AMM, AGS., MF, FC, MS, MD, RdF, RR, AV, and LFCC coordinated and carried out the bioinformatics analyses. All authors read, revised, and approved the final manuscript.

## Acknowledgments

Not applicable.

## References

1. Collette BB, Nauer CE. Scombrids of the world. An Annotated and Illustrated Catalogue of Tunas, Mackerels, Bonitos and Related Species Known to Date. FAO Species Catalogue. 1983;2:2–137. http://www.fao.org/3/ac478e/ac478e00.htm. Accessed 20 Jan 2020.

2. Hernández JJC, Ortega ATS, Castro Hernandez JJ, Santana Ortega AT. Synopsis of biological data on the chub mackerel (Scomber japonicus Houttuyn, 1782). FAO Fish Synopsis. 2000;157:1–77. https://agris.fao.org/agris-search/search.do?recordID=XF2000393177. Accessed 20 Feb 2020.

3. Velasco EM, del Arbol J, Baro J, Sobrino I. Age and growth of the Spanish chub mackerel Scomber colias off southern Spain: a comparison between samples from the NE Atlantic and the SW Mediterranean. Rev Biol Mar Oceanogr. 2011;46:27–34.

4. Gamito R, Pita C, Teixeira C, Costa MJ, Cabral HN. Trends in landings and vulnerability to climate change in different fleet components in the Portuguese coast. Fish Res. 2016;181:93–101.

5. Karakoltsidis PA, Zotos A, Constantinides SM. Composition of the commercially important mediterranean finfish, crustaceans, and molluscs. J Food Compos Anal. 1995;8:258–73.

6. Ferreira I, Gomes-Bispo A, Lourenço H, Matos J, Afonso C, Cardoso C, et al. The chemical composition and lipid profile of the chub mackerel (Scomber colias) show a strong seasonal dependence: Contribution to a nutritional evaluation. Biochimie. 2020;178:181–9.

7. Carvalho N, Perrotta RG, Isidro E. Age, growth and maturity in the chub mackerel (Scomber japonicus Houttuyn, 1782) from the Azores. Arquipélago Ciências Biológicas e Mar. 2002;19:93–9.

8. Martins MM, Skagen D, Marques V, Zwolinski J, Silva A. Changes in the abundance and spatial distribution of the Atlantic chub mackerel (Scomber colias) in the pelagic ecosystem and fisheries off Portugal. Sci Mar. 2013;77:551–63.

9. Vasconcelos J, Afonso-Dias M, Faria G. Atlantic chub mackerel (Scomber colias) spawning season, size and age at first maturity in Madeira waters. Arquipelago Life Mar Sci. 2012;29:43–51.

10. Machado AM, Felício M, Fonseca E, da Fonseca RR, Castro LFC. A resource for sustainable management: De novo assembly and annotation of the liver transcriptome of the Atlantic chub mackerel, Scomber colias. Data Br. 2018;18:276–84.

11. Catanese G, Manchado M, Infante C. Evolutionary relatedness of mackerels of the genus Scomber based on complete mitochondrial genomes: Strong support to the recognition of Atlantic Scomber colias and Pacific Scomber japonicus as distinct species. Gene. 2010;452:35–43.

12. Rodríguez-Ezpeleta N, Bradbury IR, Mendibil I, Álvarez P, Cotano U, Irigoien X. Population structure of Atlantic mackerel inferred from RAD-seq-derived SNP markers: effects of sequence clustering parameters and hierarchical SNP selection. Mol Ecol Resour. 2016;16:991–1001.

13. Ravi V, Venkatesh B. The divergent genomes of teleosts. Annu Rev Anim Biosci. 2018;6:47–68.

14. Formenti G, Theissinger K, Fernandes C, Bista I, Bombarely A, Bleidorn C, et al. The era of reference genomes in conservation genomics. Trends Ecol Evol. 2022;accepted.

15. Bolger AM, Lohse M, Usadel B. Trimmomatic: A flexible trimmer for Illumina sequence data. Bioinformatics. 2014;30:2114–20.

16. Ranallo-Benavidez TR, Jaron KS, Schatz MC. GenomeScope 2.0 and Smudgeplot for reference-free profiling of polyploid genomes. Nat Commun. 2020;11:1–10.

17. Marçais G, Kingsford C. A fast, lock-free approach for efficient parallel counting of occurrences of k-mers. Bioinformatics. 2011;27:764–70.

18. Jin JJ, Yu W Bin, Yang JB, Song Y, Depamphilis CW, Yi TS, et al. GetOrganelle: A fast and versatile toolkit for accurate de novo assembly of organelle genomes. Genome Biol. 2020;21:241.

19. Cheng H, Concepcion GT, Feng X, Zhang H, Li H. Haplotype-resolved de novo assembly using phased assembly graphs with hifiasm. Nat Methods. 2021;18:170–5.

20. Wick RR, Judd LM, Gorrie CL, Holt KE. Unicycler: Resolving bacterial genome assemblies from short and long sequencing reads. PLOS Comput Biol. 2017;13:e1005595.

21. Meng G, Li Y, Yang C, Liu S. MitoZ: A toolkit for animal mitochondrial genome assembly, annotation and visualization. Nucleic Acids Res. 2019;47:63.

22. Clavijo BJ, Garcia Accinelli G, Wright J, Heavens D, Barr K, Yanes L, et al. W2RAP: A pipeline for high quality, robust assemblies of large complex genomes from short read data. bioRxiv. 2017;:110999. doi:10.1101/110999.

23. Mapleson D, Accinelli GG, Kettleborough G, Wright J, Clavijo BJ. KAT: A K-mer analysis toolkit to quality control NGS datasets and genome assemblies. Bioinformatics. 2017;33:574–6.

24. Nurk S, Walenz BP, Rhie A, Vollger MR, Logsdon GA, Grothe R, et al. HiCanu: Accurate assembly of segmental duplications, satellites, and allelic variants from high-fidelity long reads. Genome Res. 2020;30:1291–305.

25. Wick RR, Schultz MB, Zobel J, Holt KE. Bandage: interactive visualization of de novo genome assemblies. Bioinformatics. 2015;31:3350–2.

26. Manni M, Berkeley MR, Seppey M, Simão FA, Zdobnov EM. BUSCO Update: Novel and Streamlined Workflows along with Broader and Deeper Phylogenetic Coverage for Scoring of Eukaryotic, Prokaryotic, and Viral Genomes. Mol Biol Evol. 2021;38:4647–54.

27. Gurevich A, Saveliev V, Vyahhi N, Tesler G. QUAST: Quality assessment tool for genome assemblies. Bioinformatics. 2013;29:1072–5.

28. Guan D, McCarthy SA, Wood J, Howe K, Wang Y, Durbin R. Identifying and removing haplotypic duplication in primary genome assemblies. Bioinformatics. 2020;36:2896–8.

29. Jones S, Taylor G, Chan S, Warren R, Hammond S, Bilobram S, et al. The Genome of the Beluga Whale (Delphinapterus leucas). Genes (Basel). 2017;8:378.

30. Taylor GA, Kirk H, Coombe L, Jackman SD, Chu J, Tse K, et al. The Genome of the North American Brown Bear or Grizzly: Ursus arctos ssp. horribilis. Genes (Basel). 2018;9:598.

31. Warren RL, Yang C, Vandervalk BP, Behsaz B, Lagman A, Jones SJM, et al. LINKS: Scalable, alignment-free scaffolding of draft genomes with long reads. Gigascience. 2015;4:35.

32. Warren RL. RAILS and Cobbler: Scaffolding and automated finishing of draft genomes using long DNA sequences. J Open Source Softw. 2016;1:116.

33. Li H, Durbin R. Fast and accurate long-read alignment with Burrows-Wheeler transform. Bioinformatics. 2010;26:589–95.

34. Li H. Minimap2: pairwise alignment for nucleotide sequences. Bioinformatics. 2018;34:3094–100.

35. Kim D, Paggi JM, Park C, Bennett C, Salzberg SL. Graph-based genome alignment and genotyping with HISAT2 and HISAT-genotype. Nat Biotechnol. 2019;37:907–15.

36. Kim D, Langmead B, Salzberg SL. HISAT: A fast spliced aligner with low memory requirements. Nat Methods. 2015;12:357–60.

37. Rhie A, Walenz BP, Koren S, Phillippy AM. Merqury: Reference-free quality, completeness, and phasing assessment for genome assemblies. Genome Biol. 2020;21:245.

38. Chen N. Using Repeat Masker to identify repetitive elements in genomic sequences. Curr Protoc Bioinforma. 2004;5:4–10.

39. Smit AFA, Hubley R. RepeatModeler Open-1.0. http://www.repeatmasker.org.

40. Hubley R, Finn RD, Clements J, Eddy SR, Jones TA, Bao W, et al. The Dfam database of repetitive DNA families. Nucleic Acids Res. 2016;44:D81–9.

41. Bao W, Kojima KK, Kohany O. Repbase Update, a database of repetitive elements in eukaryotic genomes. Mob DNA. 2015;6:1–6.

42. Hoff KJ, Lange S, Lomsadze A, Borodovsky M, Stanke M. BRAKER1: Unsupervised RNA-Seq-based genome annotation with GeneMark-ET and AUGUSTUS. Bioinformatics. 2015;32:767–9.

43. Hoff KJ, Lomsadze A, Borodovsky M, Stanke M. Whole-genome annotation with BRAKER. Methods Mol Biol. 2019;1962:65–95.

44. Brůna T, Hoff KJ, Lomsadze A, Stanke M, Borodovsky M. BRAKER2: automatic eukaryotic genome annotation with GeneMark-EP+ and AUGUSTUS supported by a protein database. NAR Genomics Bioinforma. 2021;3:lqaa108.

45. Li H, Handsaker B, Wysoker A, Fennell T, Ruan J, Homer N, et al. The Sequence Alignment/Map format and SAMtools. Bioinformatics. 2009;25:2078–9.

46. O’Leary NA, Wright MW, Brister JR, Ciufo S, Haddad D, McVeigh R, et al. Reference sequence (RefSeq) database at NCBI: Current status, taxonomic expansion, and functional annotation. Nucleic Acids Res. 2016;44:D733–45.

47. Yates AD, Achuthan P, Akanni W, Allen J, Allen J, Alvarez-Jarreta J, et al. Ensembl 2020. Nucleic Acids Res. 2020;48:D682–8.

48. Jones P, Binns D, Chang HY, Fraser M, Li W, McAnulla C, et al. InterProScan 5: Genome-scale protein function classification. Bioinformatics. 2014;30:1236–40.

49. Bateman A, Martin MJ, Orchard S, Magrane M, Agivetova R, Ahmad S, et al. UniProt: The universal protein knowledgebase in 2021. Nucleic Acids Res. 2021;49:D480–9.

50. Buchfink B, Xie C, Huson DH. Fast and sensitive protein alignment using DIAMOND. Nature Methods. 2014;12:59–60.

51. Buels R, Yao E, Diesh CM, Hayes RD, Munoz-Torres M, Helt G, et al. JBrowse: A dynamic web platform for genome visualization and analysis. Genome Biol. 2016;17:66.

52. Gremme G, Steinbiss S, Kurtz S. Genome tools: A comprehensive software library for efficient processing of structured genome annotations. IEEE/ACM Trans Comput Biol Bioinforma. 2013;10:645–56.

53. Li H. Tabix: Fast retrieval of sequence features from generic TAB-delimited files. Bioinformatics. 2011;27:718–9.

54. Zhang Z, Schwartz S, Wagner L, Miller W. A greedy algorithm for aligning DNA sequences. J Comput Biol. 2000;7:203–14.

55. Emms DM, Kelly S. OrthoFinder: solving fundamental biases in whole genome comparisons dramatically improves orthogroup inference accuracy. Genome Biol. 2015;16:157.

56. Edgar RC. MUSCLE: multiple sequence alignment with high accuracy and high throughput. Nucleic Acids Res. 2004;32:1792–7.

57. Capella-Gutierrez S, Silla-Martinez JM, Gabaldon T, Capella-Gutiérrez S, Silla-Martínez JM, Gabaldón T. trimAl: A tool for automated alignment trimming in large-scale phylogenetic analyses. Bioinformatics. 2009;25:1972–3.

58. Nguyen L-T, Schmidt HA, von Haeseler A, Minh BQ. IQ-TREE: A Fast and Effective Stochastic Algorithm for Estimating Maximum-Likelihood Phylogenies. Mol Biol Evol. 2015;32:268–74.

59. Kalyaanamoorthy S, Minh BQ, Wong TKF, Von Haeseler A, Jermiin LS. ModelFinder: Fast model selection for accurate phylogenetic estimates. Nat Methods. 2017;14:587–9.

60. Chapman B, Chang J. Biopython. ACM SIGBIO Newsl. 2000;20:15–9.

61. Fonseca E, Machado AM, Vilas-Arrondo N, Gomes-dos-Santos A, Veríssimo A, Esteves P, et al. Cartilaginous fishes offer unique insights into the evolution of the nuclear receptor gene repertoire in gnathostomes. Gen Comp Endocrinol. 2020;295:113527.

62. Eide M, Zhang X, Karlsen OA, Goldstone J V., Stegeman J, Jonassen I, et al. The chemical defensome of five model teleost fish. Sci Rep. 2021;11:1–13.

63. Li H, Durbin R. Inference of human population history from individual whole-genome sequences. Nature. 2011;475:493–6.

64. Barth JMI, Damerau M, Matschiner M, Jentoft S, Hanel R. Genomic Differentiation and Demographic Histories of Atlantic and Indo-Pacific Yellowfin Tuna (Thunnus albacares) Populations. Genome Biol Evol. 2017;9:1084–98.

65. Martins MM. Growth variability in Atlantic mackerel (Scomber scombrus) and Spanish mackerel (Scomber japonicus) off Portugal. ICES J Mar Sci. 2007;64:1785–90.

66. Satoh TP, Miya M, Mabuchi K, Nishida M. Structure and variation of the mitochondrial genome of fishes. BMC Genomics. 2016;17:719.

67. Hughes LC, Ortí G, Huang Y, Sun Y, Baldwin CC, Thompson AW, et al. Comprehensive phylogeny of ray-finned fishes (Actinopterygii) based on transcriptomic and genomic data. Proc Natl Acad Sci. 2018;115:6249–54.

68. Rhie A, McCarthy SA, Fedrigo O, Damas J, Formenti G, Koren S, et al. Towards complete and error-free genome assemblies of all vertebrate species. Nature. 2021;592:737–46.

69. Howe K, Chow W, Collins J, Pelan S, Pointon DL, Sims Y, et al. Significantly improving the quality of genome assemblies through curation. GigaScience. 2021;10:1–9.

70. Suda A, Nishiki I, Iwasaki Y, Matsuura A, Akita T, Suzuki N, et al. Improvement of the Pacific bluefin tuna (Thunnus orientalis) reference genome and development of male-specific DNA markers. Sci Rep. 2019;9:1–12.

71. Santos MM, Ruivo R, Capitão A, Fonseca E, Castro LFC. Identifying the gaps: Resources and perspectives on the use of nuclear receptor based-assays to improve hazard assessment of emerging contaminants. J Hazard Mater. 2018;358:508–11.

72. Bertrand S, Thisse B, Tavares R, Sachs L, Chaumot A, Bardet PL, et al. Unexpected novel relational links uncovered by extensive developmental profiling of nuclear receptor expression. PLoS Genet. 2007;3:2085–100.

73. Eide M, Rydbeck H, Tørresen OK, Lille-Langøy R, Puntervoll P, Goldstone J V., et al. Independent losses of a xenobiotic receptor across teleost evolution. Sci Rep. 2018;8:1–13.

74. López MD, Alcocer MU, Jaimes PD. Phylogeography and historical demography of the Pacific Sierra mackerel (Scomberomorus sierra) in the Eastern Pacific. BMC Genet. 2010;11:34.

75. Moura AE, Van Rensburg CJ, Pilot M, Tehrani A, Best PB, Thornton M, et al. Killer whale nuclear genome and mtDNA reveal widespread population bottleneck during the last glacial maximum. Mol Biol Evol. 2014;31:1121–31.

